## Supplemental Table for "SNX10 functions as a modulator of piecemeal mitophagy and mitochondrial bioenergetics"

| Vectors | Description | Source |
| --- | --- | --- |
| Gateway cloning vectors | | |
| pENTR1A | Entry vector | Invitrogen |
| pENTR2B | Entry vector | Invitrogen |
| pENTR3C | Entry vector | Invitrogen |
| Other vectors | | |
| pcDNA5/FRT/TO | Genomic integration vector for tetracycline-inducible expression of proteins in mammalian cells; CMV promoter. | Invitrogen |
| pCMV-VSV-G | Envelope protein for producing lentiviral and MuLV retroviral particles. | Addgene |
| pLVX-SV40-mScarlet-I-ATG16L1 | Lentiviral transfer plasmid. Based on pLVX-tight-Puro with P_tight_ promoter exchanged for SV40-mScarlet-I-ATG16L1-BGH-poly expression cassette. | This study |
| psPAX2 | 2nd generation lentiviral packaging plasmid. | Addgene |
| pMRX-IB-pSu9-HaloTag7-mGFP | Stable expression of pSu9-HaloTag7-mGFP in mammalian cells to measure mitophagy activity | Addgene |
| Plasmids made by cloning or site-directed mutagenesis | | |
| pENTR3C-SNX10 | Human SNX10 cloned into an entry vector | This study |
| pENTR3C-SNX10(Y32S) | Human SNX10 subcloned from pENTR3C-SNX10 using site-directed mutagenesis (T94A_A95G). Corresponds to variant dbSNP:rs771038257  5’-GCTATTAGTATGAATACATATCTCACTGTCAATGTAAGAATGCCAGAAGTCC-3’ and 5’-GGACTTCTGGCATTCTTACATTGACAGTGAGATATGTATTCATACTAATAGC-3’ | This study |
| pENTR3C-SNX10(R51P) | Human SNX10 subcloned from pENTR3C-SNX10 using site-directed mutagenesis (G152C).  5’-ACAATGAAAACATCCTGTGTACCAAGAAGATATAGAGAATTCGTG-3’ and  5’-CACGAATTCTCTATATCTTCTTGGTACACAGGATGTTTTCATTGT-3’ | This study |
| pENTR3C-SNX10(R51Q) | Human SNX10 subcloned from pENTR3C-SNX10 using site-directed mutagenesis (G152A). Corresponds to variant dbSNP:rs39812301  5’-ACACGAATTCTCTATATCTTCTTTGTACACAGGATGTTTTCATTGTA-3’ and  5’-TACAATGAAAACATCCTGTGTACAAAGAAGATATAGAGAATTCGTGT-3’ | This study |
| pcDNA5/FRT/TO-SNX10-EGFP | Human EGFP-tagged SNX10 subcloned from entry vectors into a pcDNA5/FRT/TO vector using Gibson Assembly. | This study |
| pcDNA5/FRT/TO-SNX10(R16L)-EGFP | Human EGFP-tagged SNX10(R16L) subcloned from entry vectors into a pcDNA5/FRT/TO vector using Gibson Assembly. | This study |
| pcDNA5/FRT/TO-SNX10(Y32S)-EGFP | Human EGFP-tagged SNX10(R51P) subcloned from entry vectors into a pcDNA5/FRT/TO vector using Gibson Assembly. | This study |
| pcDNA5/FRT/TO-SNX10(R51P)-EGFP | Human EGFP-tagged SNX10(Y32S) subcloned from entry vectors into a pcDNA5/FRT/TO vector using Gibson Assembly. | This study |
| pcDNA5/FRT/TO-SNX10(R51Q)-EGFP | Human EGFP-tagged SNX10(R51Q) subcloned from entry vectors into a pcDNA5/FRT/TO vector using Gibson Assembly. | This study |
| pcDNA5/FRT/TO-MLS-EGFP-mCherry | Double-tagged human NIPSNAP cloned into a pcDNA5/FRT/TO. | (Princely Abudu et al. 2019) |
| pLVX-SV40-mScarlet-I-Rab4A | Human mScarlet-tagged Rab4A subcloned instead of ATG16L1 into a pLVX-SV40-ATG16L1 vector. | (Pankiv et al. 2024) |
| pLVX-SV40-mScarlet-I-Rab5A | Human mScarlet-tagged Rab5A subcloned instead of ATG16L1 into a pLVX-SV40-ATG16L1 vector. | (Pankiv et al. 2024) |
| pLVX-SV40-mScarlet-I-Rab6A | Human mScarlet-tagged Rab6A subcloned instead of ATG16L1 into a pLVX-SV40-ATG16L1 vector. | (Pankiv et al. 2024) |
| pLVX-SV40-mScarlet-I-Rab7A | Human mScarlet-tagged Rab7A subcloned instead of ATG16L1 into a pLVX-SV40-ATG16L1 vector. | (Pankiv et al. 2024) |
| pLVX-SV40-mScarlet-I-Rab9A | Human mScarlet-tagged Rab9A subcloned instead of ATG16L1 into a pLVX-SV40-ATG16L1 vector. | (Pankiv et al. 2024) |
| pLVX-SV40-mScarlet-I-Rab11A | Human mScarlet-tagged Rab11A subcloned instead of ATG16L1 into a pLVX-SV40-ATG16L1 vector. | (Pankiv et al. 2024) |
| pLVX-SV40-mScarlet-I-Rab43A | Human mScarlet-tagged Rab43A subcloned instead of ATG16L1 into a pLVX-SV40-ATG16L1 vector. | (Pankiv et al. 2024) |
