## Supplemental figures and legends for "SNX10 functions as a modulator of piecemeal mitophagy and mitochondrial bioenergetics"

**Supplementary Data for Trachsel-Moncho et al.**

**Supplementary Figure 1**

**A)** Fluorescence imaging of U2OS cell lines stably expressing doxycycline-inducible SNX10-EGFP WT or the indicated ARO-linked mutants, acquired at 20x magnification using a Zeiss Axio Observer widefield microscope (Zen Blue 2.3, Zeiss). Corresponding brightfield images are displayed below the fluorescence images. Pink arrows correspond to observed vacuoles. Scale bar = 20 µm. **B)** U2OS cells stably expressing SNX10-EGFP, were treated with LysoTracker Red prior fixation. Scale bars: 10 µm. Insets: 8.40x8.40 µm. **C)** U2OS cells with stable inducible expression of SNX10-EGFP were infected with lentiviral particles to express mScarlet-RAB4/RAB5/RAB6/RAB7/RAB9/RAB11/RAB43. Nuclei were stained with Hoechst. Scale bars: 10 µm. Insets: 10.92x10.92 µm. **D)** U2OS cells with stable inducible expression of SNX10-EGFP were fixed and stained against endogenous clathrin. Nuclei were stained with Hoechst. Scale bar: 10 µm. Insets: 9.48x9.48 µm.

**Supplementary Figure 2**

**A)** U2OS cells with stable inducible expression of SNX10-EGFP were pre-treated with doxycycline for 16 hrs before the addition of DFP for 24 hrs. The cells were fixed and stained with antibodies against mitochondrial proteins for subsequent analysis. Scale bars: 10 µm. Insets: 8.57x8.57 µm **B)** Representative images of U2OS cells transfected with 20 nM siRNA: siCtrl (control) and two different siSNX10 oligoes (siSNX10 #1 and siSNX10 #2). Cells were stained with an anti-TOMM20 antibody after fixation. Images were acquired using a Nikon CREST X-Light V3 spinning disk microscope utilising a 60x oil objective. Scale bar: 10 µm. C) Quantification of the data shown in B), performed using CellProfiler software. The graph displays the area occupied by TOMM20 per cell (n =3, >100 cells per condition in each replicate). Significance was assessed by ordinary one-way ANOVA followed by Tukey's multiple comparison test. Data distribution was assumed to be normal but was not formally tested. **D)** U20S cells were reverse transfected with siSNX10 (20 nM) for 72 hrs. The cells were lysed in the well and the RNA was extracted prior to cDNA synthesis. The graph shows the difference in expression levels upon KD of the different proteins (mean values ± SEM). The values were normalized to TBP using the $2^{-\Delta\Delta Ct}$ method and then compared to siCtrl control. Significance was determined from n = 2 independent experiments, by one-way ANOVA followed by Dunnetts's multiple comparison test. Data distribution was assumed to be normal but was not formally tested. **E)** Expression levels of COX-IV were measured in control (siCtrl) and SNX10 knockdowns (siSXN10#1, siSXN10#2) upon treatment of MG132 and or DFP, across three independent experiments. Band densities were normalized to the housekeeping gene actin. Data are presented as mean ± SEM. Statistical analysis was performed using one-way ANOVA followed by Šídák's multiple comparisons test to compare each knockdown group to the control group. Data distribution was assumed to be normal but was not formally tested. * = $p < 0.05$, ** = $p < 0.01$, *** = $p < 0.001$, **** = $p < 0.0001$ , non-significant differences are not depicted.

**Supplementary Figure 3**

**A)** U2OS cells stably expressing MLS-GFP-mCherry were reverse transfected with Ctrl, SNX10 or ULK1 siRNAs (20 nM) for 72 hrs. DFP was added for the last 24 hrs and 50nM BafA1 was added 16h before fixation. Scale bar: 20 µm. The graph represents the mitolysosome area per cell from more than 1000 cells based on images taken with ImageXpress Micro Confocal (Molecular devices) at 20X magnification. The bars show the means normalized to the control (siCtrl) cells ± SEM (n=3). Significance was determined by two-way ANOVA followed by Tukey's multiple comparison test. **B)** U2OS cells stably expressing MLS-GFP-mCherry and non-tagged Parkin were reverse transfected with Ctrl and SNX10 siRNAs (20 nM) for 72 hrs. CCCP was added for the last 24 hrs and 50 nM BafA1 was added 2h before fixation. Scale bar: 20µm. The graph represents the mitolysosome area per cell from more than 1000 cells based on images taken with ImageXpress Micro Confocal (Molecular devices) at 20X magnification. The bars show the means normalized to the control (siCtrl) cells ± SEM (n=3). Significance was determined by two-way ANOVA followed by Tukey's multiple comparison test. **C)** Representative images of U2OS cells transfected with 20 nM siRNA: siCtrl (control) and two different siSNX10 oligoes (siSNX10 #1 and siSNX10 #2). Cells were stained with an anti-PDH and anti-TOMM20 antibody after fixation. Images were acquired using a Nikon CREST X-Light V3 spinning disk microscope utilising a 60x oil objective. Scale bar: 10 µm. **D)** Quantification of the data shown in C), performed using CellProfiler software. The graph displays the number of MDVs per cell, calculated as vesicles positive for TOMM20-only or PDH-only (n =3, >100 cells per condition in each replicate). Significance was assessed by two-way ANOVA followed by Dunnett's multiple comparison test. Data distribution was assumed to be normal but was not formally tested.

**Supplementary Figure 4**

**A)** Dorsal view of spatial expression pattern of snx10a and snx10b at 2, 4 and 5 dpf as demonstrated by whole-mount in situ hybridization (WM-ISH) using an internal antisense - (AS) probe. Scale bar = 200 µm. Images are representative of 3 experiments. Control 2 dpf larvae hybridized to a sense probe (S). **B)** Illustration of sgRNA binding regions on *snx10a* and *snx10b* gene respectively. **C)** Temporal expression levels of *bnip3* and *bnip3l* transcripts in DMSO and DMOG treated WT zebrafish larvae at 3 dpf. The graph shows the fold change in transcript levels relative to β-actin and normalized to DMSO $2^{-\Delta\Delta Ct}$ levels. Error bars indicate mean ± SEM. Data is collected from 3 individual experiments. Significance was determined by two-way ANOVA test to compare all groups with the 2 variables. **D)** Temporal expression levels of *cox-iv* and *samm50* transcripts in control and *snx10ab*_DKO zebrafish larvae treated with or without 100 µm DMOG at 3 dpf. The graph shows the fold change in transcript levels relative to β-actin and normalized to DMSO $2^{-\Delta\Delta Ct}$ levels. Error bars indicate mean ± SEM. Data is collected from 3 individual experiments. Significance was determined by one-way ANOVA test to compare all groups with the individual variable. Data distribution was assumed to be normal but was not formally tested. * = $p < 0.05$, ** = $p < 0.01$, *** = $p < 0.001$, **** = $p < 0.0001$ , non-significant differences are not depicted. **E)** Representative dot plots showing the region selected for FACS analysis from control and snx10ab DKO zebrafish larvae at 3 dpf using MitoSOX reagent. H2O2 was added to the water for 1 hr as a positive control. **F)** Representative FACS plot showing oxidative stress in control, and snx10ab DKO zebrafish larvae at 3 dpf using the MitoSOX reagent. H2O2 added in water served as positive control.

**Supplementary Table 1**

Table S1 summarizes the plasmids utilized throughout the study, detailing their names and respective descriptions.

**Movie 1**

**Time-lapse fluorescence imaging of SNX10-EGFP-expressing U2OS cells treated with MitoTracker Red.** This movie depicts live imaging of U2OS cells with inducible expression of SNX10-EGFP and treated with MitoTracker Red 30 min prior to imaging. The acquisition lasted 2 min at intervals of 500 milliseconds, and the playback rate is 6 frames per second (fps). Scale bar = 10 µm. Related to Fig. 3C.

#### Supplementary Figure 1

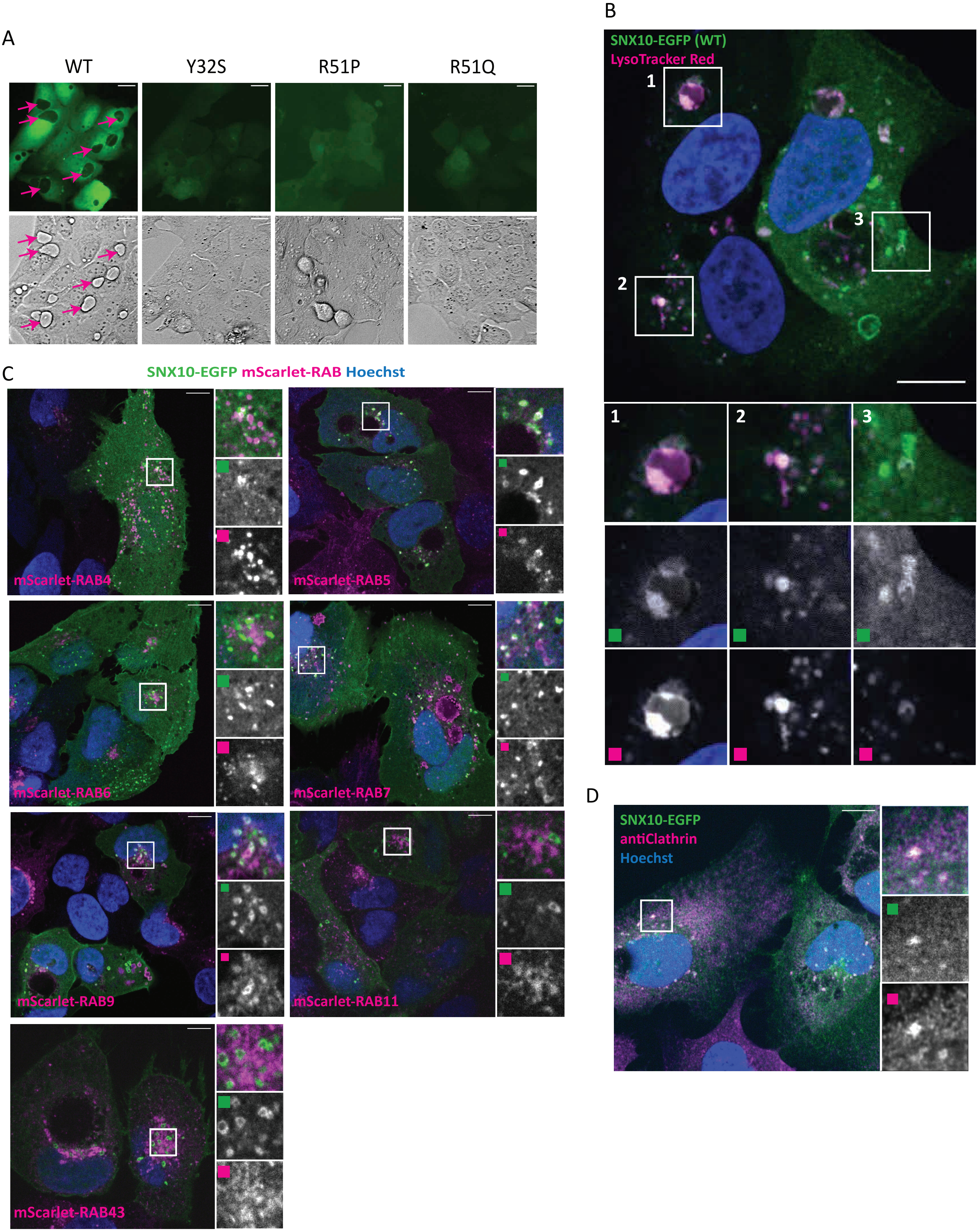

### Supplementary Figure 2

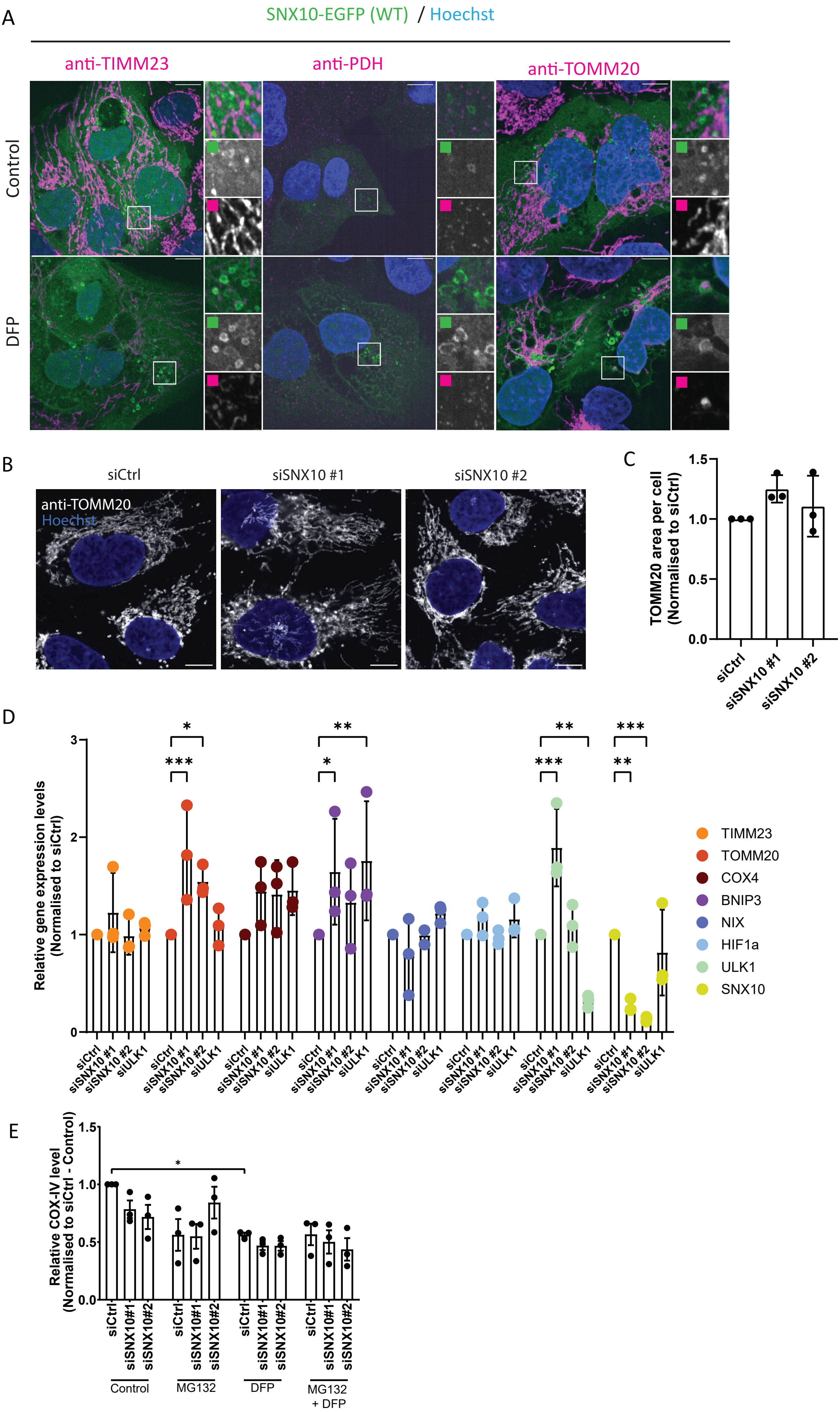

### Supplementary Figure 3

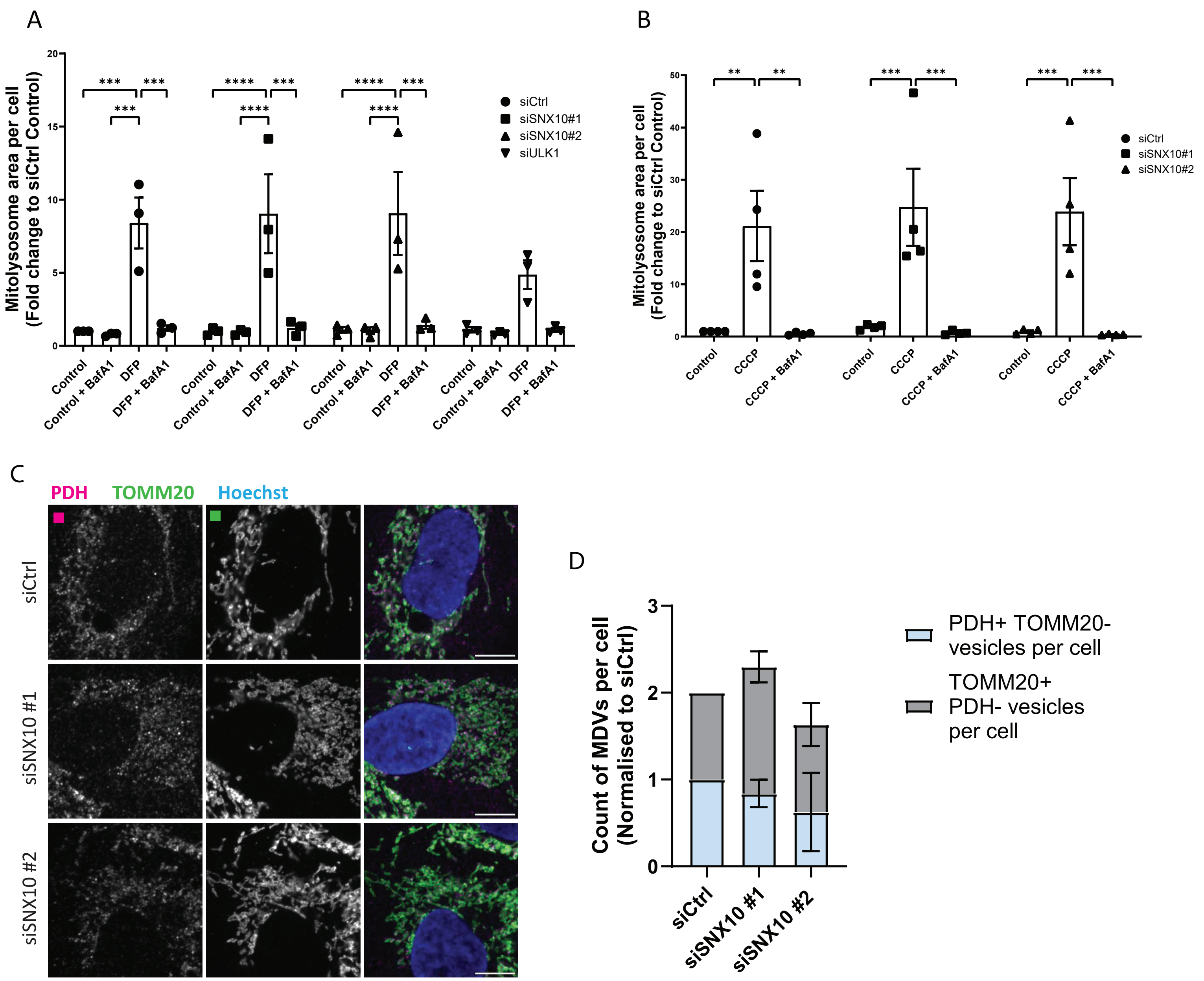

### Supplementary Figure 4

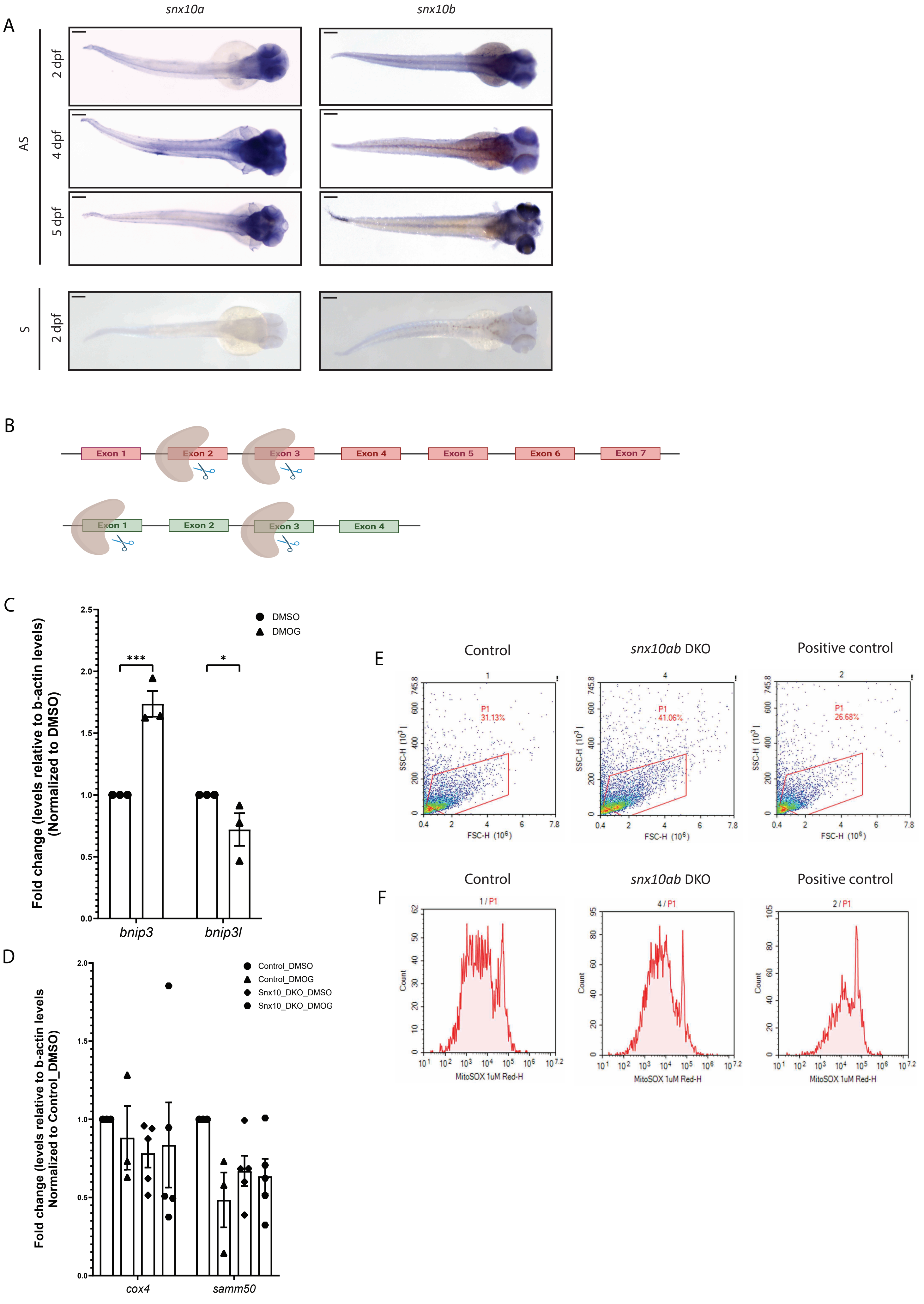
